# A conserved epilepsy-associated gene co-expression module identifies increased metabolic rate as a shared pathomechanism

**DOI:** 10.1101/2025.02.03.636180

**Authors:** Jingyi Long, Spencer G. Jones, Ana Serna, Boyd van Reijmersdal, Franziska Kampshoff, Sara Aibar, Patrik Verstreken, Martijn A. Huynen, Kevin Lüthy, Mireia Coll-Tané, Annette Schenck

**Affiliations:** Department of Human Genetics, Radboud University Medical Center, 6525 GA, Nijmegen, the Netherlands; Donders Institute for Brain, Cognition and Behaviour, Radboud University Medical Center, 6525 EN, Nijmegen, the Netherlands; VIB-KU Leuven Center for Brain & Disease Research, 3000 Leuven, Belgium; KU Leuven, Department of Human Genetics, 3000 Leuven, Belgium; VIB-KU Leuven Center for Brain & Disease Research, 3000 Leuven, Belgium; KU Leuven, Department of Neurosciences, Leuven Brain Institute, 3000 Leuven, Belgium; Department of Medical Biosciences, Radboud University Medical Center, 6525 GA, Nijmegen, the Netherlands; Institute of Human Genetics, UK Essen, University of Duisburg-Essen School of Medicine, 45147 Essen, Germany

**Keywords:** **Keyword** Epilepsy, gene co-expression network, metabolic rate, AMPK, *Drosophila*

## Abstract

Epilepsy is a mechanistically complex, incompletely understood neurological disorder. To uncover novel converging mechanisms in epilepsy, we used *Drosophila* whole-brain single-cell RNA-sequencing to refine and characterize a previously proposed human epilepsy-associated gene co-expression network (GCN). We identified a conserved co-expressed module of 26 genes, which comprises fly orthologs of 13 epilepsy-associated genes and integrates synaptic and metabolic functions. Over one-third of the *Drosophila* pan-neuronal knockdown models targeting this module exhibited altered seizure-like behaviors in response to mechanical or heat stress. These recapitulated seizures associated with four epilepsy-associated genes, identified two novel epilepsy candidate genes, and three genes knockdown of which conferred seizure protection. Most knockdown models with altered seizure susceptibility showed changes in metabolic rate and levels of phosphorylated adenosine monophosphate-activated protein kinase (AMPK), a key regulator of cellular energy homeostasis. Enhancing AMPK activity increased seizure resistance in a dose-dependent manner. Our findings show that *Drosophila* single-cell expression data and behavior can aid functional validation of human GCNs and highlight a role for metabolism in modifying seizure susceptibility.

**SUMMARY STATEMENT:** Integrating *Drosophila* single-cell RNA-sequencing data with seizure-like behavior and metabolic rate assays, we functionally characterized a human epilepsy-associated gene network, revealing metabolic regulation as a critical factor underlying seizure susceptibilities.

## INTRODUCTION

Epilepsy, characterized by the spontaneous recurrence of unprovoked seizures, impacts over 50 million people globally and manifests in combination with a large number of neurological, cognitive, and psychosocial consequences (Fisher et al., 2014, Thijs et al., 2019). Despite recent advances, epilepsy treatment remains challenging, with about one-third of patients showing resistance to current anti-epileptic drugs (Mesraoua et al., 2023). This emphasizes the need to explore the biological underpinnings of epilepsy to enable the development of novel treatment strategies. Next-generation sequencing technologies have markedly accelerated the identification of genetic factors implicated in epilepsy and suggested that genetic predisposition contributes to up to 80% of epilepsy cases (Hildebrand et al., 2013, Thakran et al., 2020) *De novo* mutations in more than 1500 genes have been linked to epilepsy (Zhang et al., 2023), yet their underlying pathogenic mechanisms and how they contribute to epileptogenesis remain to be determined.

Systems biology and network analyses are potent methods for exploring the molecular processes and pathways involved in diseases (Parikshak et al., 2015). Gene co-expression network (GCN) analysis is useful for identifying clusters of genes that show similar expression patterns under various conditions and thus likely share biological functions (Van Dam et al., 2018). This approach has revealed a first pro-convulsant gene network and proposed Sestrin 3 as a regulator among diverse epilepsies (Johnson et al., 2015). Nonetheless, validating the multitude of genes within GCNs remains a significant challenge, as experimentation is both costly and time-intensive (Van Dam et al., 2018). Delahaye-Duriez and colleagues used genome-wide gene expression data from previously published post-mortem healthy human brains (Ramasamy et al., 2014) to perform Weighted Gene Co-expression Network Analysis (WGCNA) and Differential Co-expression Analysis (DiffCoEx). They identified a co-expression network of 320 genes (termed M30 network) that is enriched in genes associated with mono- and polygenic epilepsy, providing opportunities for epilepsy treatment exploration (Delahaye-Duriez et al., 2016). They also reported concerted (dys)regulation of the network in several epilepsy-related datasets, but experimental investigation of potential convergent mechanisms was not attempted.

*Drosophila melanogaster* is an excellent model to study the etiology of epilepsy due to its efficiency, cost-effectiveness, and ease of genetic manipulation. Assays to induce seizure-like behaviors are well-established, and 81% of human epilepsy genes have orthologs in this organism (Fischer et al., 2023). The availability of comprehensive single-cell brain expression datasets in *Drosophila* (Davie et al., 2018) provided a novel opportunity to functionally characterize GCNs associated with epilepsy. Using these data, we identified M30 genes with conserved neuronal co-expression in this evolutionarily distant organism and isolated a highly co-expressed module of 26 genes for in-depth examination. Genetic manipulation of this fly co-expression module induced seizure-like behavior in several models, identifying both known epilepsy-associated genes as well as novel candidate genes and modifiers. It also revealed strong functional coherence of the module linked to regulating metabolic rate, implicating a novel converging neuronal mechanism in a specific group of genetic epilepsies.

## RESULTS

### Identification of a conserved epilepsy-associated gene co-expression module in *Drosophila*

To provide novel insights into the genetic causes of epilepsy and reveal shared mechanisms, we integrated a previously identified gene co-expression network enriched in epilepsy-associated genes generated based on human post-mortem brain tissue bulk RNA sequencing (RNA-seq) data (the M30 network, comprising 320 genes) (Delahaye-Duriez et al., 2016) with single-cell gene expression data in *Drosophila* (Davie et al., 2018) (**Fig. 1**).

**Figure 1.**
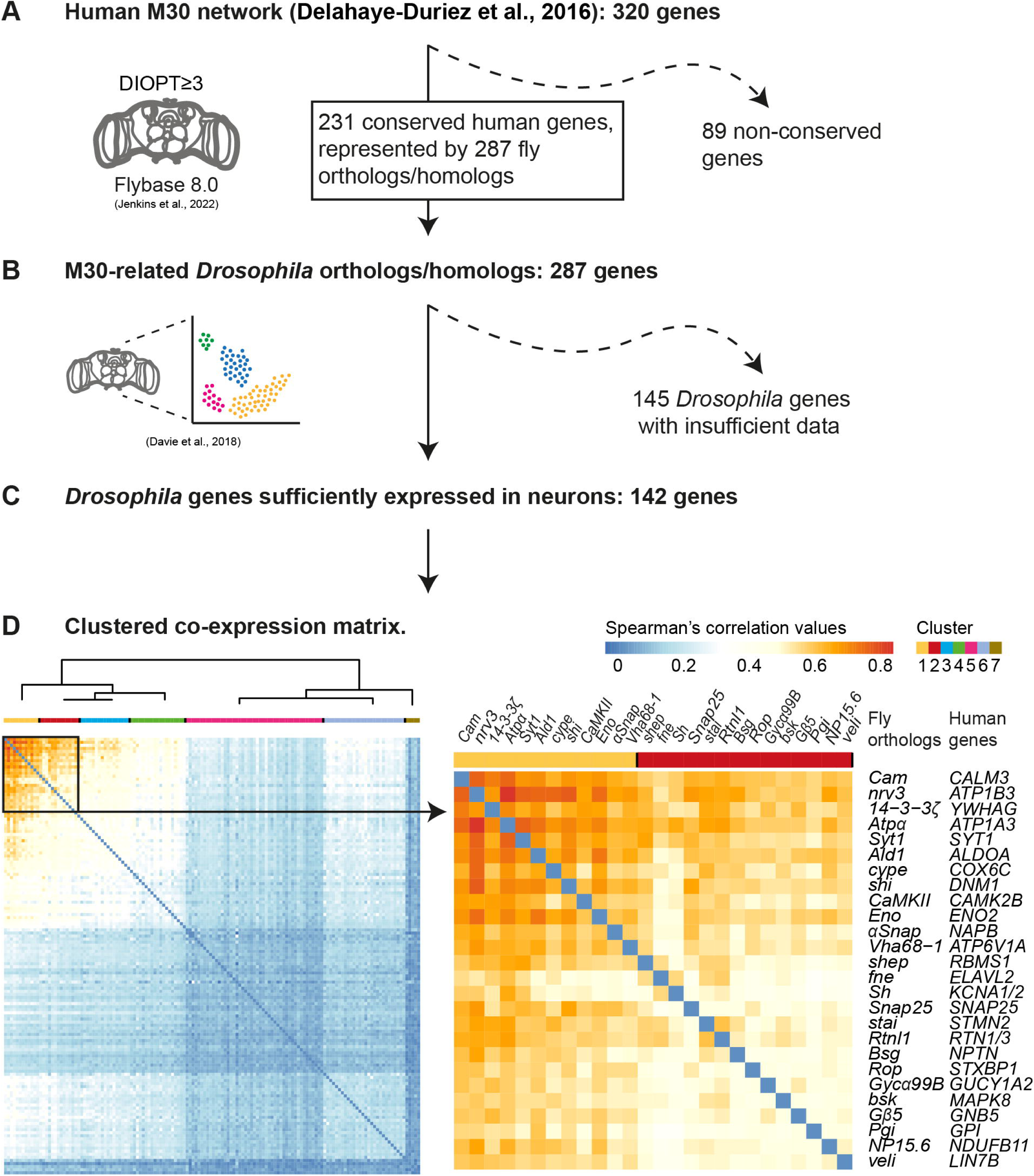
A conserved neuronal co-expression module of 26 *Drosophila* genes based on the human epilepsy-associated M30 network and whole-brain *Drosophila* scRNA-seq atlas. Workflow diagram representing the steps from the human epilepsy-associated M30 gene network to the identification of the highly co-expressed genes 26 genes. **(A)** Annotation of M30 *Drosophila* orthologs. **(B)** Extracting genes with >5000 normalized scRNA-seq counts. **(C)** Calculations of the pairwise Spearman’s *ρ* values and clustering. **(D)** Left: The hierarchically clustered heatmap represents the co-expression of 142 *Drosophila* genes. Right: It highlights a module of 26 highly co-expressed fly genes (indicated with a black square, magnified on the right). Each line or column represents a unique gene. The color indicates co-expression values. To the right of the 26 x 26 gene matrix, the fly orthologs of the human M30 genes are indicated. The 26 gene module is composed of 2 clusters: 12 most co-expressed genes in the core cluster (top left, cluster 1 in yellow) and 14 further genes in cluster 2 (in red).

First, we used the DRSC (*Drosophila* RNAi Screening Center) Integrative Ortholog Prediction Tool (DIOPT) scores (Hu et al., 2021) to determine the conservation and *Drosophila* orthologs of the 320 human genes of the M30 network (**Table S1,** see Materials and Methods). We identified 287 M30-related *Drosophila* genes, encompassing one-to-one, many-to-one (a sole fly gene representing more than one human gene), and one-to-many (more than one fly gene with a similar DIOPT score representing a human gene) orthologs (**Fig. 1A**, **Table S2**). To determine the co-expression of these genes in *Drosophila*, we retrieved the whole-brain single-cell RNA sequencing (scRNA-seq) data from Davie et al. (Davie et al., 2018) (**Fig. 1B**). Pairwise Spearman’s correlation values were calculated for the 142 *Drosophila* genes with sufficient expression across most neuronal subtypes present in the dataset (see Materials and Methods) (**Fig. 1C**, genes listed in **Table S3**, Spearman’s *ρ* values (r_s_) in **Table S4**). To unravel which genes show the highest degree of expression similarity and, hence, are most likely functionally related, hierarchical clustering was performed and represented in a heatmap (**Fig. 1D**). Visual inspection identified a highly co-expressed module of 26 genes, distinct from the background of lowly-correlated genes. These 26 genes belonged to two hierarchical clusters. The first ‘core’ cluster included the 12 genes with the highest co-expression values (0. 52 < r_s_ < 0.84). The second cluster of 14 genes was characterized by r_s_ values ranging between 0.36 and 0.61 within this cluster, and r_s_ values up to 0.68 with genes in the top-scoring core cluster.

Multiple genes in both co-expressed clusters are associated with epilepsy (**Table 1**), as classified in OMIM (Online Mendelian Inheritance in Man; <u>OMIM</u>) and documented in PubMed (nih.gov). In summary, we identified an epilepsy-associated module comprising 26 genes characterized by evolutionary conserved co-expression.

**Table 1.**
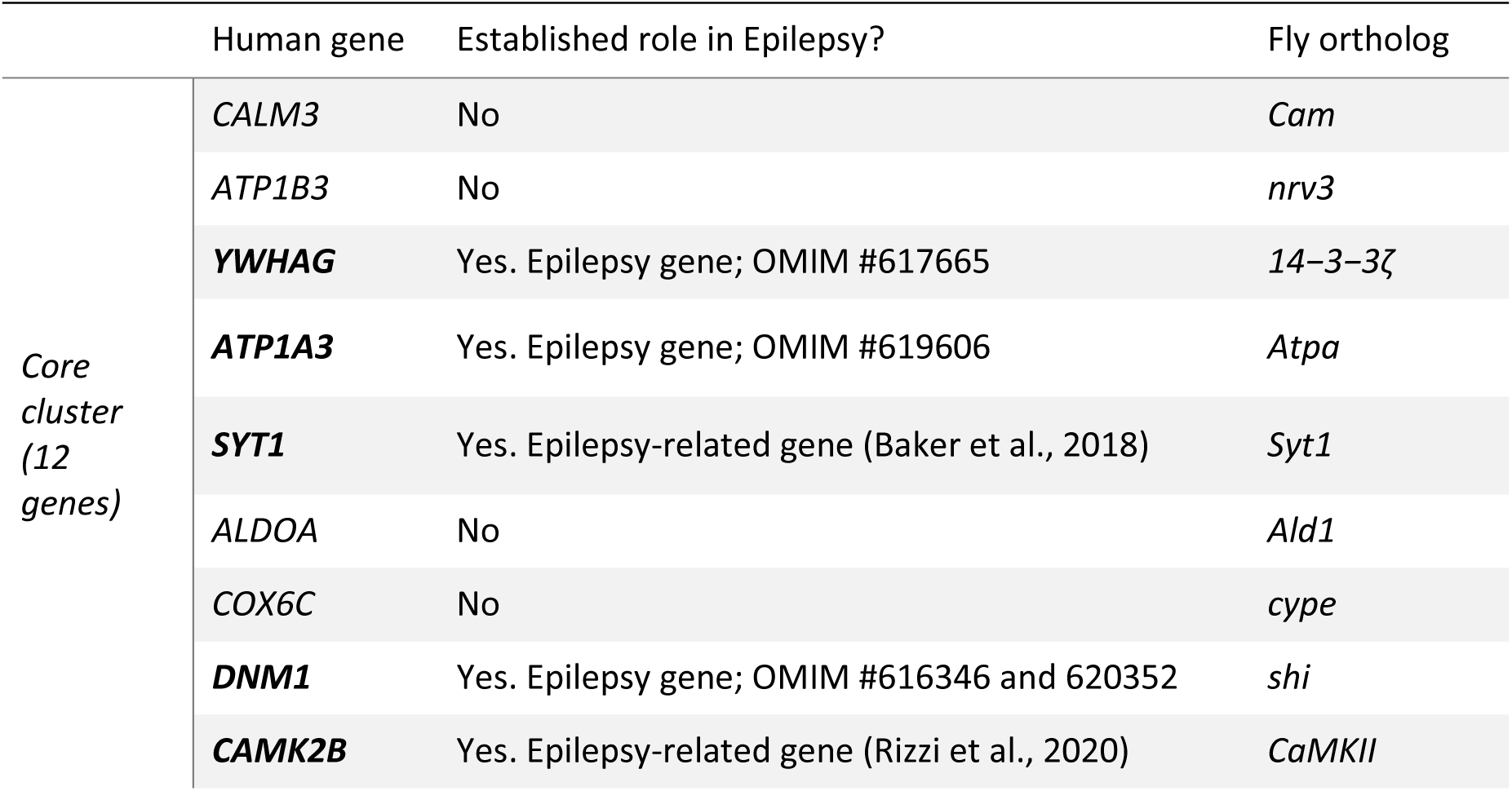
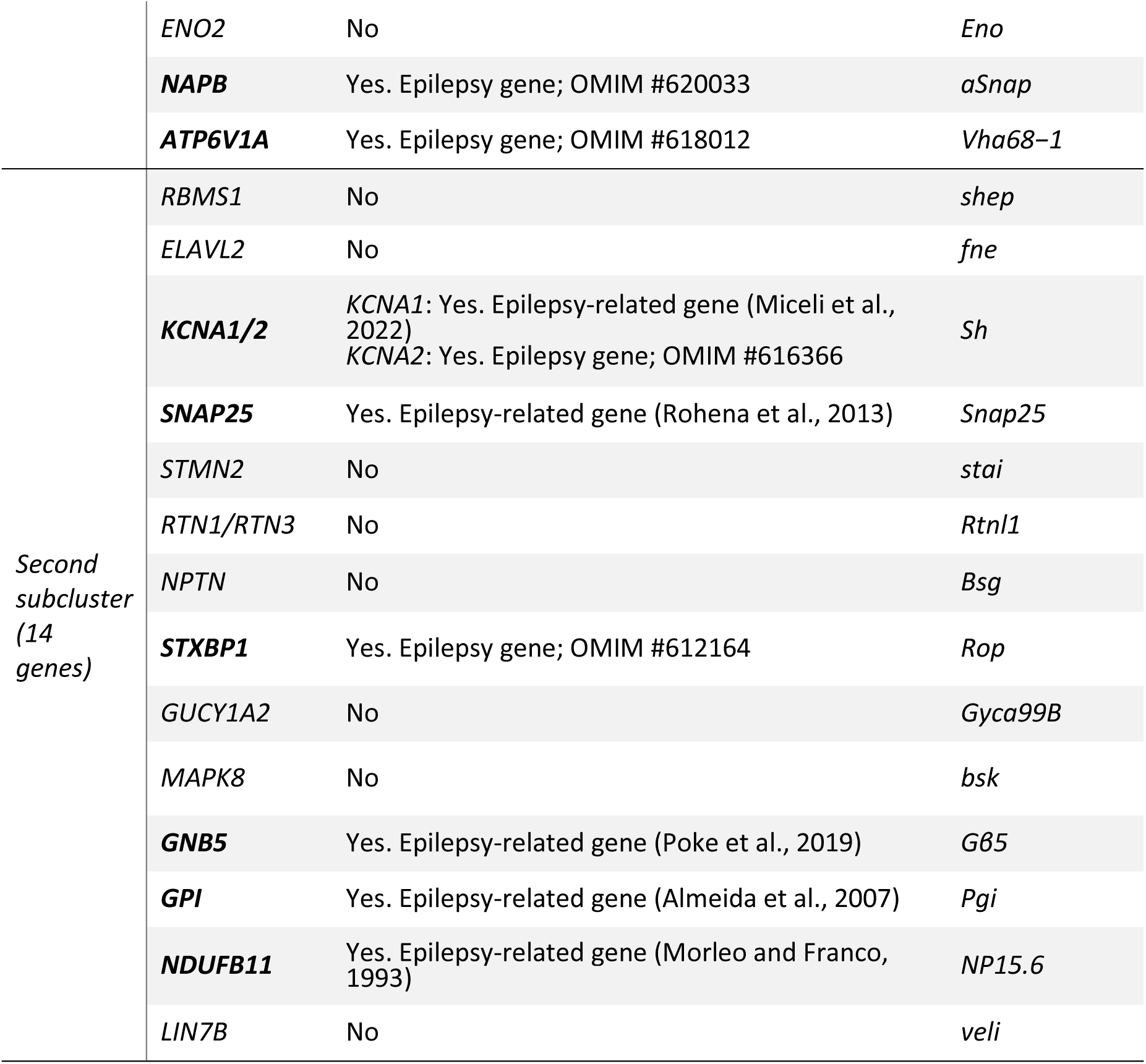
Catalog of the 26 genes comprised in the highly co-expressed module. Human genes, their currently established implication in epilepsy, and their fly orthologs are indicated. Bold gene names highlight both established epilepsy genes and epilepsy-related genes. Epilepsy genes, identified in the Gene-Phenotype Relationship tables of OMIM as genes with epilepsy or epilepsy-related terms (e.g., epileptic encephalopathy). Epilepsy-related genes were identified as those for which epilepsy or seizures are listed in OMIM’s clinical features section or found case reports using terms like “epilepsy” or “seizure”. Abbreviations: OMIM - Online Mendelian Inheritance in Man.

### The epilepsy-associated and highly conserved co-expression module links genes with synaptic and metabolic functions

As a first step toward characterizing the molecular nature of the 26-gene module we identified, we performed Kyoto Encyclopedia of Genes and Genomes (KEGG) pathway and gene ontology (GO) enrichment analysis comparing this module against the genome-wide background using g:Profiler (Raudvere et al., 2019). Whereas the human M30 cluster was highly enriched in genes relevant to neural processes, such as synaptic transmission, synaptic vesicle transport, and gamma-aminobutyric acid (GABA) signaling (Delahaye-Duriez et al., 2016) [confirmed by our recent query (**Fig. 2A, top**)], three of the four significantly enriched KEGG pathways of our co-expression module were related to carbohydrate metabolic processes (**Fig. 2A, bottom**). Similarly, GO-term analysis for biological processes associated with our module ranked carbohydrate-related metabolic processes among the top-scored ontology terms [e.g., ‘glucose homeostasis’ (FDR = 1.82×10^-3^, fold enrichment = 28.69), ‘ATP metabolic process’, (FDR = 4.96×10^-4^, fold enrichment = 27.66); ‘carbohydrate metabolic homeostasis’, (FDR = 0.047, fold enrichment = 5.41)], in addition to synaptic and other neuronal processes (**Fig. 2B**). We also identified other enriched functions among the 26 gene module, neither identified by Delahaye-Duriez and colleagues (Delahaye-Duriez et al., 2016) nor us when performing GO-term analysis of the human M30 genes. These include G-protein-coupled receptor signaling and cognitive functions such as learning and memory (**Fig. 2B**, **Table S5**). In total, 449 significantly enriched GO terms were identified for the co-expressed *Drosophila* 26 gene module versus 367 for the human M30 cluster. Of these, only 155 overlapped between the human and *Drosophila* GO terms (**Fig. 2C**). Whereas this overlap is significant, as expected from a smaller list selected from a larger one, the large divergence is remarkable. In particular, the increased number of significantly enriched GO terms despite the lower number of genes (and hence potential statistical power) illustrates that our approach to identifying evolutionary conserved, biologically coherent clusters within the M30 cluster worked.

**Figure 2.**
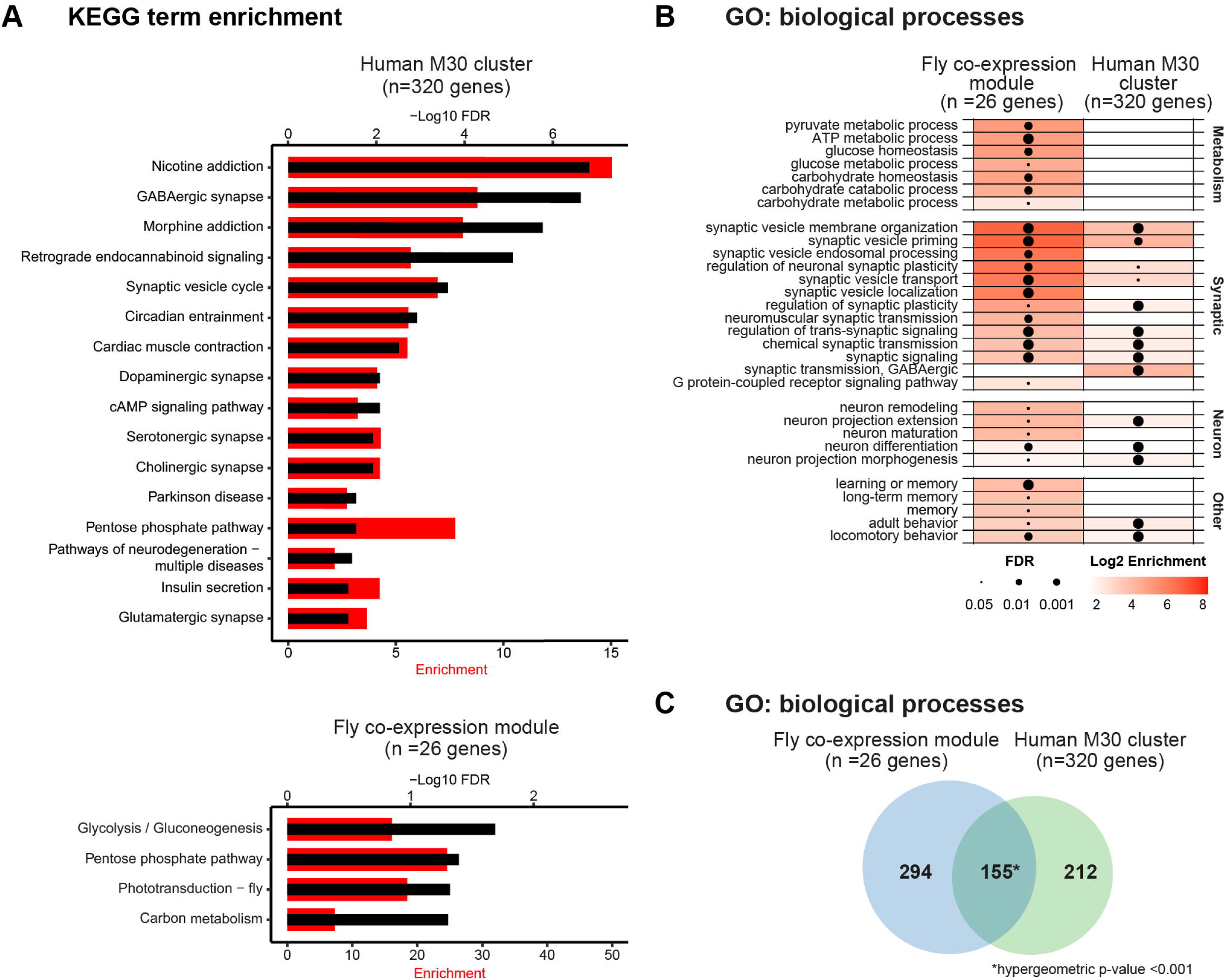
The 26 gene co-expression module links genes with synaptic and metabolic functions. **(A)** KEGG pathway analysis of the human M30 cluster (top) and the *Drosophila* co-expression module (bottom). The fly co-expression module’s enriched KEGG pathways primarily relate to carbohydrate metabolic processes, whereas these are not enriched in the human M30 cluster. For each KEGG pathway, the bars show the -Log10 FDR (black, axis on the top) and the fold enrichment (red, axis on the bottom). **(B)** Enriched GO biological processes of the *Drosophila* co-expression module and the human M30 cluster. Only significantly enriched, representative GO biological processes (FDR < 0.05) are shown. A full list of significant terms is shown in supplemental Table S5. The size of the dots represents the FDR, and the color intensity of each grid represents the Log2 enrichment. The carbohydrate metabolism process ranked highly in the GO-term analysis of biological processes. **(C)** Venn diagram illustrating the overlap of GO biological processes that are enriched in our *Drosophila* co-expression module compared with the human M30 cluster. The asterisk indicates significance: * *p* < 0.001. Abbreviations: FDR – False Discovery Rate

Finally, we evaluated the contribution of epilepsy-associated genes and their gene orthologs to the M30 and our 26-gene module. According to current knowledge (OMIM and PubMed, see Material and Methods), there were 31 epilepsy genes, and 22 epilepsy-related genes among the 320 human genes in the M30 cluster (**Table S6**), together corresponding to 17% (53/320 genes). Among the *Drosophila* 26 genes are 7 epilepsy genes and 6 epilepsy-related genes, together summing up to 50% (13/26 genes) and a statistically significant 3-fold enrichment in epilepsy-associated genes compared to the human M30 cluster (*p*-value = 3.45×10^-5^). This suggests that the functional themes identified in our module are highly relevant to epilepsy.

### More than a third of the neuronal knockdown fly models in the co-expression gene module have altered seizure susceptibility

To address whether and which of the identified co-expressed *Drosophila* genes can be implicated in seizure-like behaviors, we took advantage of the UAS-Gal4 system (Brand and Perrimon, 1993) and the *nSyb-Gal4* driver to induce pan-neuronal knockdown models by RNA interference (RNAi) (Dietzl et al., 2007, Perkins et al., 2015). We systematically targeted the core cluster, encompassing the 12 genes with the highest co-expression and seven genes in the second interrelated cluster, which we selected based on high co-expression values, molecular and biological functions (**Tables S4 and S5**), and available tools. In total, we used 44 RNAi lines (from four RNAi libraries) to knock down 19 genes, with at least two independent constructs per gene whenever available. In parallel, we crossed the *nSyb-Gal4* driver to the corresponding genetic background control lines of each of the RNAi lines. We found that pan-neuronal knockdown of *nrv3* (RNAi-1, −2, and −3), *Atpα* (RNAi-1 and 2), *Cam* (RNAi-3), *Ald1* (RNAi-3), *Eno* (RNAi-1 and 2), *shi* (RNAi-2), *cype* (RNAi-3), *Vha68-1* (RNAi-3), *ɑSnap* (RNAi-3), *shep* (RNAi-3), and *Rtnl1* (RNAi-2) led to developmental lethality (**Table S7**), which precluded further evaluation of seizure-like behavior.

Seizure susceptibility in adult flies can be assessed upon exposure to mechanical (bang-sensitive) or heat stress, which are believed to be caused by different underlying molecular mechanisms (Fischer et al., 2023, Mituzaite et al., 2021). We evaluated both types of seizure-like behaviors in the adult viable RNAi models. In the mechanically induced seizure-like behavior assay, flies are placed in a laboratory vortex at the highest speed to hyperstimulate sensory inputs and assess the resulting seizure-like behavior (Kuebler and Tanouye, 2000) (**Fig. 3A**). Seizure-like behavior is reflected by uncontrolled wing and leg movements while flies lay on their backs or by paralysis (Kuebler and Tanouye, 2000, Lasko and Lüthy, 2021). Pan-neuronal knockdown of *14-3-3ζ* (RNAi-1 and −2), *CaMKII* (RNAi-1 and −2), *cype* (RNAi-1), and *NP15.6* (RNAi-1) showed significantly increased seizure frequency compared with their isogenic controls (**Fig. 3A’**). Of these four genes, the human orthologs of all but *cype* are known to be associated with epilepsy (Morleo and Franco, 1993, Ramocki et al., 2010, Rizzi et al., 2020) (**Table 1**). The increased seizure-like behavior in *Drosophila* and its presence in the epilepsy-associated highly co-expression module identify *COX6C*, the human ortholog of *cype*, as a potential epilepsy gene.

**Figure 3.**
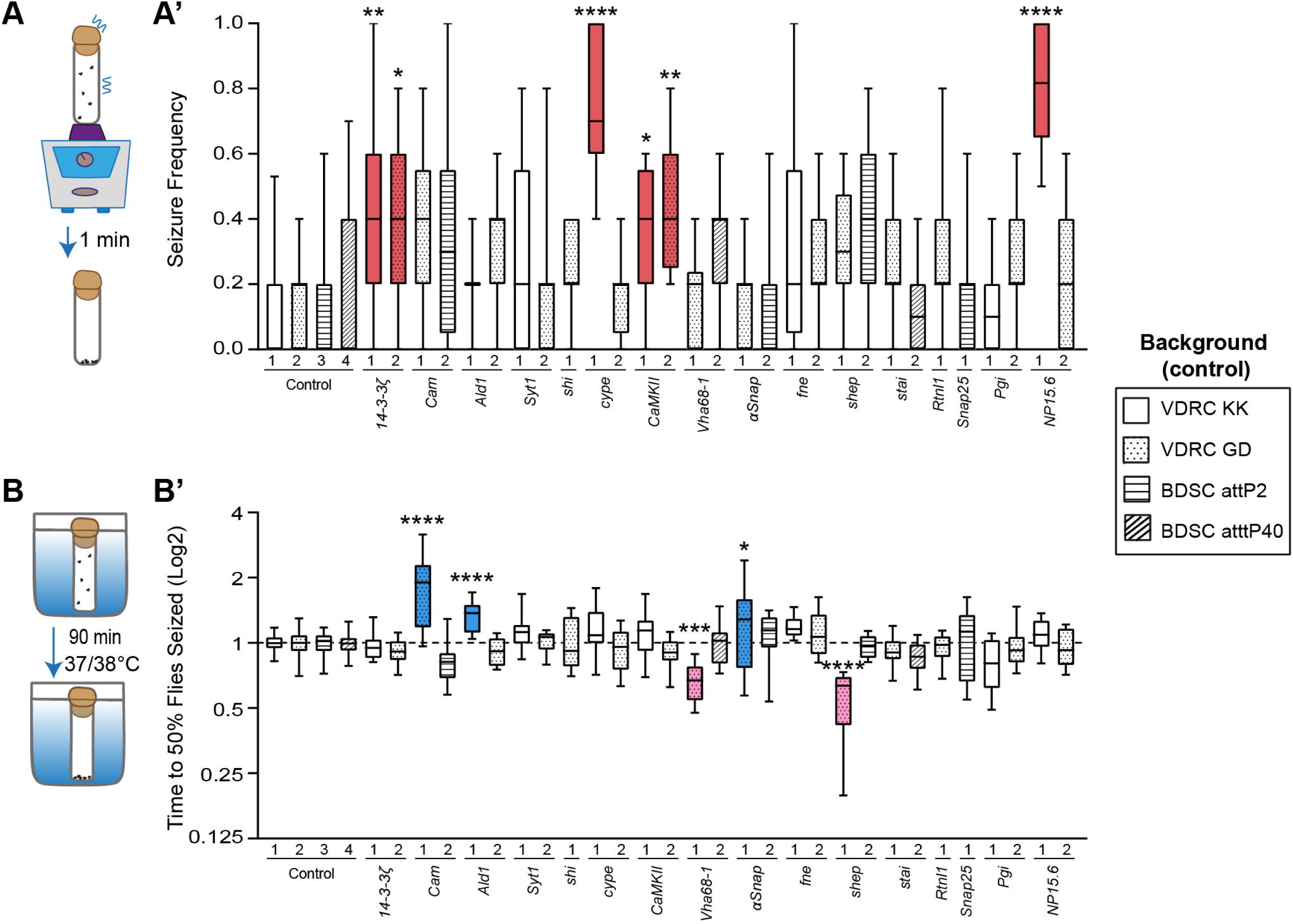
Pan-neuronal knockdown models with altered seizure susceptibility upon either mechanical or heat stress induction. **(A)** Schematic representation of the paradigm used to assess seizure-like behavior upon mechanical stress. **(A’)** Seizure frequency in all viable screened RNAi lines and isogenic background controls, crossed with pan-neuronal driver *nSyb-Gal4*. Red indicates knockdown models that have significantly higher seizure frequency. Pan-neuronal knockdown of *14-3-3ζ* (RNAi-1 and 2), *cype* (RNAi-1), *CaMKII* (RNAi-1 and 2), and *NP15.6* (RNAi-1) leads to a significant increase in seizure frequency compared to their genetic background controls (*14-3-3ζ* RNAi-1, *p* = 0.0025 and RNAi-2, *p* = 0.014; *cype* RNAi-1, *p* < 0.0001; *CaMKII* RNAi-1, *p* = 0.014 and RNAi-2, *p* = 0.0012; *NP15.6* RNAi-1, *p* < 0.0001). Kruskal-Wallis test with Dunn’s correction for multiple testing. N = 12 vials per RNAi line, 4-6 males per vial. **(B)** Schematic representation of the paradigm used to assess seizure-like behavior upon heat stress. **(B’)** Normalized average time to which 50% of the flies are seized in all viable screened RNAi lines and isogenic background controls, crossed with pan-neuronal driver *nSyb-Gal4*. Pink indicates knockdown models that show a significantly shorter time until 50% of the flies show seizure-like behavior at 37°C or 38°C compared to their genetic controls (*Vha68-1* RNAi-1, *p* = 0.0007; *shep* RNAi-1, *p* < 0.0001), indicating that they have higher heat sensitivity and are more prone to seizures. Blue indicates knockdown models that show a significant increase in the time it takes the flies to show seizure-like behavior (*Cam* RNAi-1, *p* < 0.0001; *Ald1* RNAi-1, *p* < 0.0001; *ɑSnap* RNAi-1, *p* = 0.018), indicating that they have lower heat sensitivity and are protected against seizure susceptibility. One-way ANOVA test with Šidák correction for multiple testing. N = 10-12 vials per RNAi line, 4-6 flies per vial. Data are represented as boxplots that extend from the 25^th^ and 75^th^ percentiles, with the median indicated. Whiskers indicate the 5^th^ and 95^th^ percentiles. *p*-values are indicated as follows: * *p* < 0.05, ** *p* < 0.01, *** *p* < 0.001, **** *p* < 0.0001. Abbreviations: VDRC-Vienna *Drosophila* Resource Center; BDSC - Bloomington *Drosophila* Stock Center.

Next, we investigated which of our models show seizure-like behavior upon heat stress (Burg and Wu, 2012, Mituzaite et al., 2021) (**Fig. 3B**). Flies that exhibit increased susceptibility to seizure-like behavior drop to the bottom of the vials faster than wild-type animals and show uncontrolled movements or paralysis. We found that pan-neuronal knockdown of *Vha68-1* (RNAi-1) led to a significant increase in heat-sensitive seizure-like behavior compared with their isogenic background controls (**Fig. 3B’**). Notably, its human ortholog, *ATP6V1A*, is an established epilepsy gene (Fassio et al., 2018) (**Table 1**). The *Drosophila* ortholog of *RBMS1*, *shep* (RNAi-1), also showed a strikingly increased susceptibility to seizure-like behavior (**Fig. 3B’**). Interestingly, pan-neuronal knockdown of *Cam* (RNAi-1), *Ald1* (RNAi-1), and *ɑSnap* (RNAi-1) caused decreased heat sensitivity, suggesting a protective role against heat-sensitive seizure-like behavior (**Fig. 3B’**). Taken together, our experimental approach revealed altered seizure susceptibility in more than a third (11 of 29, 38%) of the tested neuronal knockdown models within the *Drosophila* co-expression module. It effectively recapitulated the seizures of four epilepsy-associated genes, identified two novel epilepsy candidate genes, and proposed seizure-protective effects for another three genes.

### A majority of pan-neuronal knockdown models with altered seizure susceptibility have alterations in metabolic rate

The enrichment of KEGG pathways and GO terms related to carbohydrate and ATP metabolism among the identified co-expression module (**Fig. 2A-B**), along with the fact that epileptic seizures are marked by increased neuronal activity and higher ATP requirements (Rho and Boison, 2022), prompted us to investigate if changes in the metabolic state may occur more widely among the models with altered seizure susceptibility. Carbon dioxide (CO_2_) production can serve as a reliable indicator of substrate oxidation and energy expenditure, thus providing a valuable indication of the metabolic state (Van Voorhies et al., 2004). We measured CO_2_ production using respirometers that measure changes in gas volume when CO_2_ is absorbed, which causes the pressure to drop and the liquid columns in the respirometers’ capillaries to rise (Yatsenko et al., 2014) (**Fig. 4A**). First, we tested the four genes that exhibited increased seizure frequency upon mechanical stress. Pan-neuronal knockdown of *14-3-3ζ*, *CaMKII,* and *NP15.6* resulted in a significant increase in CO_2_ production compared with isogenic controls (**Fig. 4B-D**), illustrating increased metabolic rates. Pan-neuronal knockdown of *cype* did not affect the metabolic rate (**Fig. 4E**).

**Figure 4.**
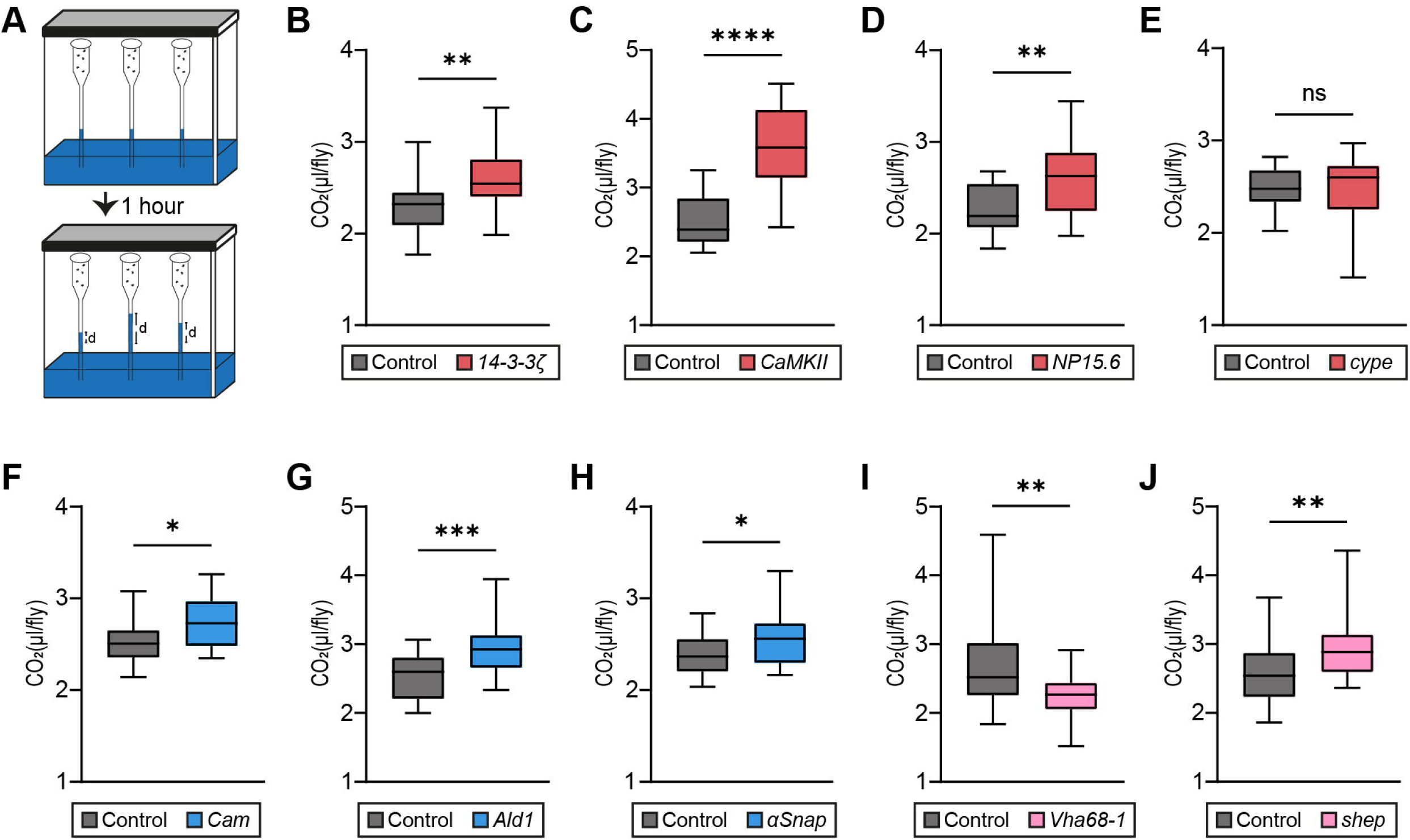
Pan-neuronal knockdown models with seizure-like behavior show altered metabolic rate. **(A)** Schematic representation of the paradigm used to assess metabolic rate by a respirometry system, used to measure CO_2_ production. Changes in gas volume after CO_2_ absorption led to a decrease in pressure and an increase in the fluid level inside the respirometer capillaries within a sealed chamber. The distance by which the liquid ascends in the micropipette after 1 hour is depicted as distance “d”. **(B-E)** CO_2_ production in the pan-neuronal knockdown models (*nSyb-Gal4 > UAS-RNAi*) with increased seizure frequency in the mechanically induced seizure assay (in red), and their isogenic background controls (genetic background of the respective RNAi line crossed with the same driver, *nSyb-Gal4/ +*). **(B-D)** Pan-neuronal knockdown of *14-3-3ζ*, *CaMKII*, and *NP15.6* leads to a significant increase in CO_2_ production and thus in their metabolic rates (*14-3-3ζ*, *p* = 0.0041; *CaMKII*, *p* < 0.0001; *NP15.6*, *p* = 0.0019). **(E)** Pan-neuronal knockdown of *cype* has no effect on metabolic rate (*p* = 0.95). **(F-H)** CO_2_ production in the pan-neuronal knockdown models with lower heat sensitivity (in blue), and their isogenic background controls. Pan-neuronal knockdown of *Cam*, *Ald1*, and *ɑSnap* lead to a significant increase in CO_2_ production and thus in their metabolic rate (*Cam*, *p* = 0.017; *Ald1*, *p* = 0.0003; *ɑSnap*, *p* = 0.026). **(I-J)** CO_2_ production in the pan-neuronal knockdown models with higher heat sensitivity (in pink), and their isogenic background controls. **(I)** Pan-neuronal knockdown of *Vha68-1* leads to lower CO_2_ production and thus a decrease in its metabolic rate (*p* = 0.0096). **(J)** Pan-neuronal knockdown of *shep* leads to an increase in CO_2_ production and thus in its metabolic rate (*p* = 0.0098). N = 18-24 groups per genotype, with five flies per group. Data are represented as boxplots that extend from the 25th and 75th percentiles, with the median indicated. Whiskers indicate the minimum and maximum. Two-tailed unpaired t-test or Mann-Whitney test, based on the normality of the distribution. *p*-values are indicated as follows: * *p* < 0.05, ** *p* < 0.01, *** *p* < 0.001, **** *p* < 0.0001.

Surprisingly, pan-neuronal knockdown of the three genes that caused lower heat sensitivity (an indication of seizure protection) − *Cam*, *Ald1*, and *ɑSnap* − also showed increased CO_2_ production (**Fig. 4F-H**). Furthermore, pan-neuronal knockdown of the two genes causing increased heat sensitivity, *Vha68-1* and *shep* (**Fig. 4I-J**), showed significantly dysregulated metabolic rate as well, albeit in opposite directions, with *Vha68-1* being the only one among the tested models exhibiting a decrease in metabolic rate (**Fig. 4I**). Collectively, our findings show that pan-neuronal knockdown models exhibiting alterations in seizure susceptibility show metabolic dysregulation. Notably, this also applies to genes not associated with the identified metabolic KEGG and GO terms, such as *shep* and *aSnap*, which have no previous reported function in energy metabolism/regulation of metabolic rate (**Table S8**).

### Neuronal models with altered seizure susceptibility show increased AMPK phosphorylation

Adenosine monophosphate-activated protein kinase (AMPK) plays a pivotal role in safeguarding cellular energy homeostasis. Cellular energy stress, such as glucose deprivation (Zhang et al., 2017) or low ATP levels (Hardie and Carling, 1997), causes AMPK activation by phosphorylation (Lin and Hardie, 2018). This, in turn, increases catabolism and decreases anabolism through the phosphorylation of key proteins in multiple pathways, including mTOR complex 1 (mTORC1), lipid homeostasis, glycolysis, and mitochondrial homeostasis (Herzig and Shaw, 2018). Therefore, we asked whether increased metabolic rates in the *Drosophila* pan-neuronal knockdown models with seizure-like behavior are associated with changes in AMPK phosphorylation. We addressed this question by subjecting head extracts of the above-investigated knockdown models to quantitative western blot analysis using anti-phospho-AMPK (p-AMPK) antibodies. This revealed that pan-neuronal knockdown of *14-3-3ζ*, *CaMKII*, *NP15.6*, *Cam*, *Ald1*, and *ɑSnap*, all characterized by increased metabolic rate, led to a significant threefold or higher increase in phosphorylated AMPK levels compared to their respective isogenic controls (**Fig. 5A-5C’ and E-G’**). Pan-neuronal knockdown of *Vha68-1,* causing heat-sensitive seizure-like behavior and a decrease in metabolic rate, and *shep,* causing heat-sensitive seizure-like behavior and an increase in metabolic rate, also resulted in a significantly but modest increase (fold change < 2) in AMPK phosphorylation (**Fig. 5H-I’**). Surprisingly, the pan-neuronal knockdown of *cype* also manifested a significant rise in AMPK phosphorylation (**Fig. 5D-D’**), despite maintaining a normal metabolic rate. Together, these results show that alterations in seizure susceptibility in most pan-neuronal knockdown models are paralleled by variable increases in AMPK phosphorylation levels.

**Figure 5.**
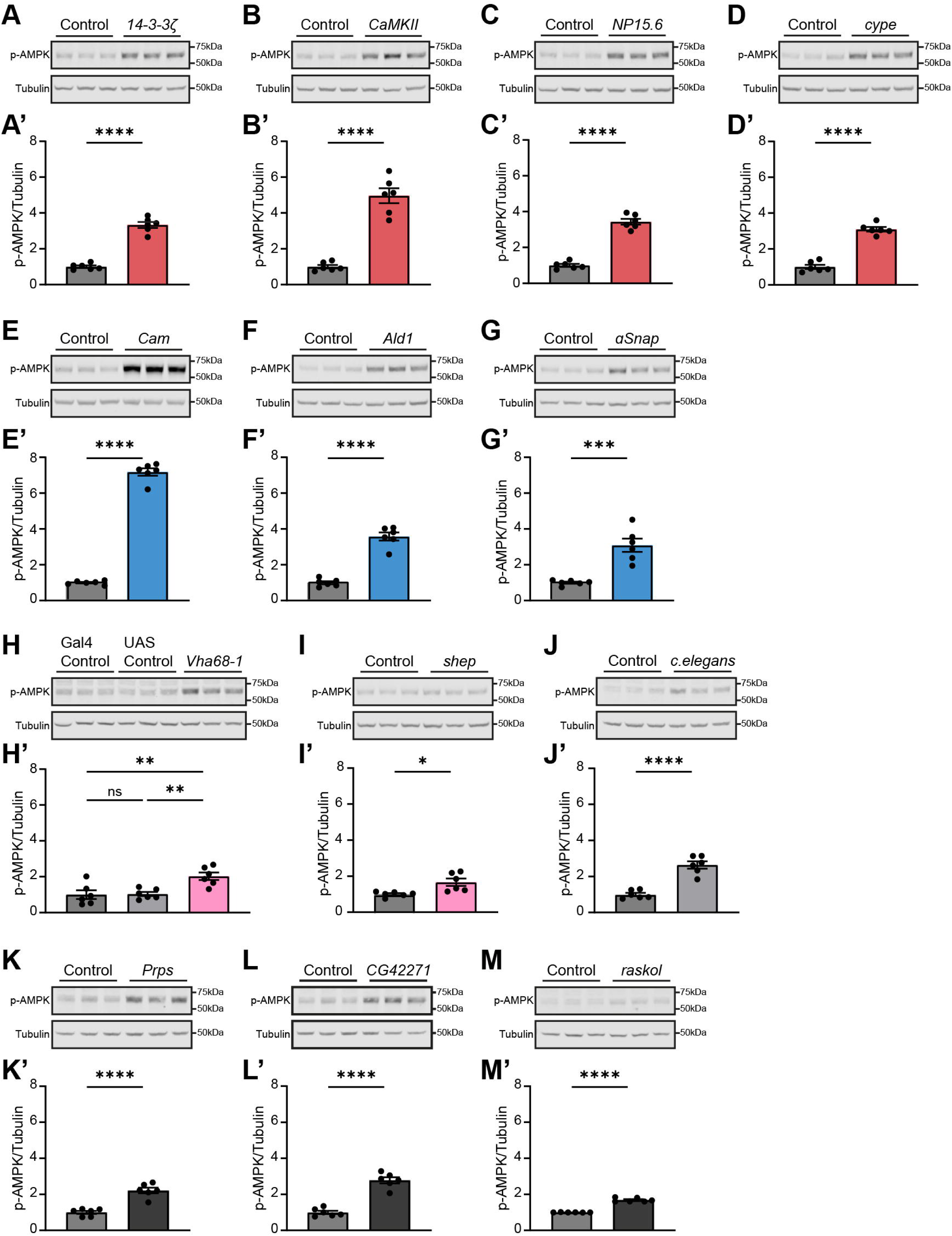
Pan-neuronal knockdown models with altered seizure susceptibility, but also negative controls with induced non-sense or unrelated RNAi, show different degrees of increased AMPK phosphorylation. **(A-M)** Representative western blots showing three biological replicates of head homogenates from adult flies containing *nSyb-Gal4* and UAS-RNAi genetic elements or their genetic background controls using anti-phospho-AMPK (p-AMPK) and anti-tubulin antibodies, and **(A’-M’)** their quantification. p-AMPK levels are normalized against the tubulin loading control. **(A-G and A’-G’)** Pan-neuronal knockdown of *14-3-3ζ*, *CaMKII*, *NP15.6*, *cype*, *Cam*, *Ald1* and *ɑSnap* leads to a significant increase in phosphorylated AMPK compared to their genetic background controls (*14-3-3ζ*, *p* < 0.0001; *CaMKII*, *p* < 0.0001; *NP15.6*, *p* < 0.0001; *cype*, *p* < 0.0001; *Cam*, *p* < 0.0001; *Ald1*, *p* < 0.0001; *ɑSnap*, *p* = 0.0002). **(H and H’)** Pan-neuronal knockdown of *Vha68-1* leads to a slight but significant increase in phosphorylated AMPK levels compared to their genetic background controls (*Vha68-1*, dark grey: *p* = 0.0064; light gray: *p* = 0.0083). Flies containing only the *UAS-Vha68-1-RNAi* construct show similar levels of phosphorylated AMPK than flies containing only *nsyb-Gal4* (*p* = 0.9989). **(I and I’)** Pan-neuronal knockdown of *shep* leads to a slight but significant increase in phosphorylated AMPK levels compared to their genetic background controls (*shep*, *p* = 0.01). **(A’-I’)** Colors according to behavioral phenotypes in epilepsy assays, as in figures 3 and 4. **(J and J’)** Pan-neuronal induction of a UAS-RNAi construct targeting a *C. elegans* gene leads to a significant increase in phosphorylated AMPK compared to its genetic background control (*p* < 0.0001). **(K-N and K’-N’)** Pan-neuronal knockdown of *Prps*, *CG42271* and *raskol* show a significant increase in phosphorylated AMPK compared to their genetic background controls (*Prps*, *p* < 0.0001; *CG42271*, *p* < 0.0001; *raskol*, *p* < 0.0001). Data are shown as mean ± SEM. **(A’-G’, I’-M’)** Two-tailed unpaired t-test. **(H’)** One-way ANOVA test with Šidák correction for multiple testing. *p*-values are indicated as follows: * *p* < 0.05, ** *p* < 0.01,*** *p* < 0.001,**** *p* < 0.0001.

To further investigate the increase in AMPK phosphorylation, which we observed in our *nSyb-Gal4 > UAS-RNAi* models without exception, we included additional controls. First, we investigated a non-specific control fly line in which *nSyb-Gal4* drives an RNAi hairpin sequence targeting a *Caenorhabditis elegans* (*C. elegans*) gene with no target in *Drosophila*. Unexpectedly, pan-neuronal induction of the *C. elegans-*specific RNAi also resulted in increased AMPK phosphorylation (**Fig. 5J-J’**). To determine whether a UAS-RNAi construct alone could affect AMPK phosphorylation without being induced, we representatively evaluated p-AMPK levels on flies carrying the *UAS-Vha68-1-RNAi* construct but not the *nSyb-Gal4* (UAS control). We found no differences in the levels of phosphorylated AMPK between this and our standard Gal4 control (**Fig. 5H-H’**). Furthermore, we tested three additional RNAi lines targeting orthologs of the human M30 cluster, which are not included in the 26 gene co-expression module. Pan-neuronal knockdown of *Prps*, *CG42271* and *raskol* all resulted in increased AMPK phosphorylation (**Fig. 5K-M’**). Combined, these data argue that activation of the RNAi machinery itself appears to affect AMPK phosphorylation, regardless of the targeted gene. However, whereas pan-neuronal knockdown of the M30 orthologs not associated with our module (*Prps*, *CG42271*, *raskol*, the *C. elegans*-specific RNAi), as well as *Vha68.1* and *shep* knockdown resulted in a modest increase in phosphorylated AMPK levels by 1.69 - 2.78 fold, pan-neuronal knockdown of *14-3-3ζ*, *NP15.6*, *Ald1,* and above all *CaMKII* and *Cam* strongly elevated levels of p-AMPK (fold change compared to isogenic controls: 3.33 - 7.17), suggesting that these genes regulate AMPK phosphorylation specifically.

### AMPK regulates seizure susceptibility

To address the causality of increased AMPK phosphorylation levels, questioned by at least some degree of unspecific activation upon RNAi, for seizure-like behavior, we aimed to determine whether neuronal manipulation of AMPK in flies alters their seizure susceptibility. We pan-neuronally overexpressed either *UAS-AMPKα^M^*, *UAS-AMPKα^T184D^*, or *UAS-AMPKα^K57A^*, a wild-type, a constitutively active, and a dominant-negative form of the kinase, respectively (Johnson et al., 2010, Swick et al., 2013). In the mechanically induced seizure-like behavior assay, the expression of neither *UAS-AMPKα^M^, UAS-AMPKα^T184D^*, nor *UAS-AMPKα^K57A^* had any effect on seizure frequency (**Fig. 6A**). In contrast, in the heat induced seizure-like behavior assay, the expression of the constitutively active AMPK (*UAS-AMPKα^T184D^*) showed a slight but significantly reduction in heat sensitivity (longer time to seizure) compared to isogenic controls (**Fig. 6B**). Because the efficiency of the UAS-Gal4 system is increased at higher temperatures (Brand and Perrimon, 1993), we also evaluated heat-sensitive seizure-like behavior on flies that were reared at 28 °C instead of 25 °C, and induced seizures at 42 °C instead of 38 °C. Under these stringent conditions, the reduced heat sensitivity phenotype caused by the overexpression of constitutive active AMPK (*UAS-AMPKα^T184D^*) was more pronounced, and the overexpression of wild-type AMPK (*UAS-AMPKα^M^*) also acquired the phenotype (**Fig. 6C**). Expression of the dominant-negative AMPK allele had no effect on heat-induced seizure susceptibility (*UAS-AMPKα^K57A^*; **Fig. 6B,C**). In summary, increasing AMPK levels and/or activity is sufficient to improve resistance to heat induced seizure-like behavior in a dose-dependent manner.

**Figure 6.**
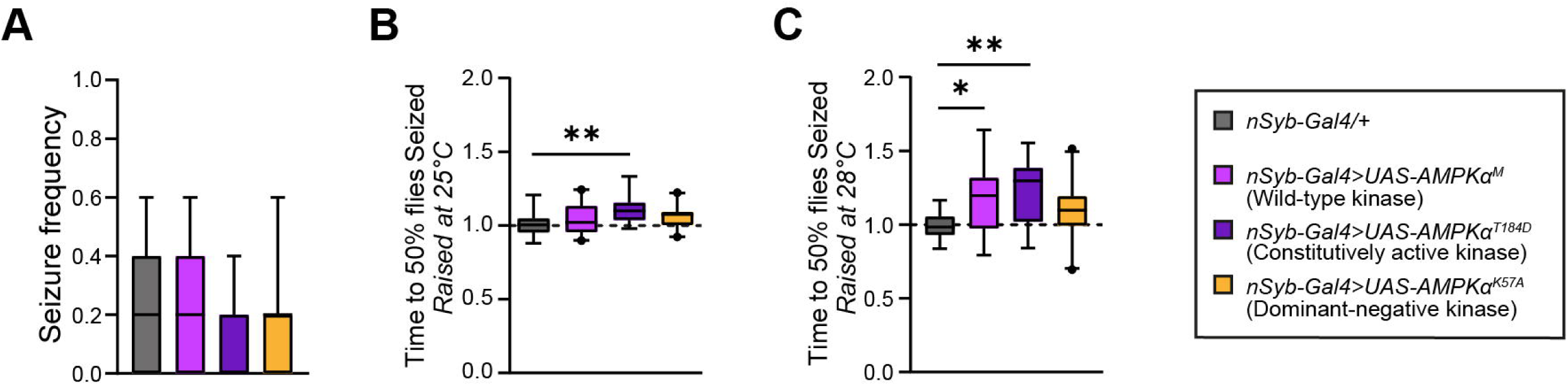
Overexpression wild-type and constitutively active AMPK alters heat induced seizure susceptibility. **(A)** Seizure frequency upon mechanical stress is not altered in flies overexpressing either wild-type (*nSyb-Gal4 > UAS-AMPKɑ^M^*), a constitutively active (*nSyb-Gal4> UAS-AMPKɑ^T184D^*), or a dominant-negative form of AMPK (*nSyb-Gal4> UAS-AMPKɑ^K57A^*) under the control of pan-neuronal driver, compared to their background control. N = 18 vials with 5 males per vial. Kruskal-Wallis test with Dunn’s correction for multiple testing. **(B)** Pan-neuronal overexpression of a constitutively activate form of AMPK leads to increased average time for 50% of the flies showing seizure-like behavior (*nSyb-Gal4 > UAS-AMPKɑ^T184D^*, *p* = 0.0012) compared to their background control, indicating that they have a lower heat sensitivity and thus a protective effect towards seizure susceptibility. N = 18-20 vials, 4-5 males per vial. One-way ANOVA test with Dunnett’s correction for multiple testing. **(C)** Pan-neuronal overexpression of wild-type and constitutive active AMPK leads to lower heat sensitivity (*nSyb-Gal4 > UAS-AMPKɑ^M^, p* = 0.023; *nSyb-Gal4 > UAS-AMPKɑ^T184D^, p* = 0.0011). Flies were raised at 28°C and exposed to a temperature of 42°C to induce seizures. N = 16-21 vials, 5 males per vial. One-way ANOVA test with Dunnett’s correction for multiple testing. Data are represented as boxplots that extend from the 25^th^ and 75^th^ percentiles, with the median indicated. Whiskers indicate the 5^th^ and 95^th^ percentiles. *p*-values are indicated as follows: * *p* < 0.05; ** *p* < 0.01.

## DISCUSSION

Research in epilepsy has evolved from focusing on single-gene investigations to studying genetic networks and functional processes. Despite significant advancements, the sheer number of genes often identified in bioinformatics approaches, such as genome-wide GCNs, poses challenges for validating hypotheses and further studying experimental phenotypes and mechanisms further. In the current study, we leveraged *Drosophila* whole-brain single-cell RNA-sequencing data to refine the previously reported M30 epilepsy gene network (Delahaye-Duriez et al., 2016). This identified a highly co-expressed cluster of 26 genes, whose composition suggested a functional connection between synaptic machinery and energy metabolism in epilepsy. Through genetic manipulation and metabolic rate measurements, we confirmed this prediction in several *Drosophila* models, uncovering a convergence of epilepsy genes operating at the synapse on the regulation of metabolic state. The molecular mechanisms underlying seizure and metabolic modulation in our module appear to still be heterogenous but include dysregulation of AMPK activity, which alone was sufficient to confer protection against heat-induced seizure-like behavior.

### Relevance of the *Drosophila* co-expression module to epilepsy

Utilizing *Drosophila* whole-brain single-cell RNA-sequencing data, we defined a highly conserved co-expression module of 26 genes, which includes fly orthologs of 13 epilepsy-associated genes (50%). Compared to 17% of epilepsy-associated genes in the human M30 cluster, this highlights an increased relevance of the *Drosophila* module to epilepsy. Moreover, our experimental models successfully recapitulated increased seizure susceptibility linked to four epilepsy-associated genes (*14-3-3ζ*, *CaMKII*, *NP15.6*, and *Vha68-1*). *YWHAG* (*14-3-3ζ* in *Drosophila*) encodes a member of the highly conserved 14-3-3 adaptor protein family that are enriched at synapses (Foote et al., 2015, Zhang and Zhou, 2018) and may be a strong predictor for cognitive impairment in Alzheimer’s disease (Oh et al., 2024 preprint). *ATP6V1A* (*Vha68-1* in *Drosophila*) encodes the A-subunit of the V1 domain of the vacuolar-ATPase (V-ATPase), essential for vesicle acidification and synaptic transmission (Bodzeta et al., 2017, Forgac, 2007). Both *YWHAG* and *ATP6V1A* are classified as epilepsy genes linked to developmental and epileptic encephalopathy (OMIM #617665 and OMIM #618012, respectively). CaMKII is a multifunctional serine/threonine kinase critical for synaptic plasticity, learning, and memory (Wang et al., 2000) and *CAMK2B* is linked to intellectual disability that can occur with co-morbid epilepsy (Kury et al., 2017) (OMIM #617799). *NDUFB11* (*NP15.6* in *Drosophila*), a component of mitochondrial complex I, is implicated in microphthalmia with linear skin defects syndrome, which includes seizures (Morleo and Franco, 1993) (OMIM #300952). Therefore, our models provide a robust platform for validating and further exploring genes involved in epilepsy.

In addition, our findings suggest a seizure-modulatory, protective effect upon the knockdown of three genes, *Ald1*, *Cam*, and *ɑSnap*. *ALDOA* (*Aldolase A*), the human ortholog of *Ald1*, is a key enzyme in the fourth step of glycolysis (Morais et al., 2017). Interestingly, while neuronal excitability has traditionally been attributed to ion channels and synaptic transmission, glucose metabolism also plays a crucial role in its regulation. Glycolysis inhibition has been shown to reduce neuronal excitability (Shao et al., 2018), highlighting the importance of metabolic pathways in modulating seizure susceptibility. While the reduced levels of CaMKII are associated with increased seizure susceptibility in our models, our results indicate that knockdown of *Cam* (Calmodulin) confers protection against seizures. Calmodulin plays a crucial role in calcium signaling by binding to Ca²⁺, which activates CaMKII (Junho et al., 2020). Cam and CaMKII can thus be expected to operate synergistically. Our finding, though seemingly contradictory, may align with research indicating that both decreased and increased activity of CaMKII signaling can cause epileptic phenotypes (Proietti Onori and Van Woerden, 2021, Rizzi et al., 2020) as well as with reports showing that calcium antagonistic effects are observed in several anticonvulsants, suggesting that calmodulin inhibition might also reduce seizures (Chaisewikul et al., 2001, Walden et al., 1992). The mechanisms underlying these discrepancies remain unclear, both in the literature and our study, but they support a mixed nature of epilepsy-predisposing and protective factors contributing to the identified co-expression module.

We found that neuronal knockdown of *Drosophila ɑSnap* confers protection against seizures. *Drosophila ɑSnap* and its human orthologs *NAPA* and *NAPB* are members of the evolutionarily conserved SNAP (soluble N-ethylmaleimide-sensitive fusion attachment proteins) family involved in vesicle docking and fusion (Andreeva et al., 2006). Mutations in *NAPB* have recently been implicated in epilepsy (Mignon-Ravix et al., 2023), and *Napb* knockout mice show seizures (Burgalossi et al., 2010), seemingly opposing the directionality of our finding. Nevertheless, no mutations in *NAPA* have been linked to epilepsy and viable hypomorphic *Napa* mutant mice do not show seizures (Chae et al., 2004). Thus, the decreased susceptibility to seizures observed in our study may result from a divergent function of the human *NAPA/B* paralogs and their two-to-one orthology in *Drosophila*. Alternatively, it is possible that seizures in individuals with mutations in *NAPB* (Mignon-Ravix et al., 2023) and knockout mice (Burgalossi et al., 2010) originate from βSnap deficiency in cells or tissues, not targeted in our neuronal knockdown approach.

Lastly, our study also identified two novel candidate epilepsy genes, based on guilt by association (Piovesan et al., 2015), through our co-expressed cluster and the obtained seizure-like phenotypes in *Drosophila*: *RBMS1* (*shep*) and *COX6C* (*cype*). *RBMS1* encodes an RNA-binding protein that may contribute to epilepsy through the misregulation of its mRNA targets, while *COX6C* encodes a subunit of complex IV in the mitochondrial respiratory chain, crucial for oxidative phosphorylation (OXPHOS) (Wang et al., 2022). The connection between epilepsy and OXPHOS function is further supported by NDUFB11 (NP15.6), a gene also in our cluster, associated with seizure-like behavior in our model as well as with seizures in patients (Morleo and Franco, 1993). Since mitochondrial OXPHOS deficiency accelerates glycolysis (Zheng, 2012), this aligns with our observed protective effect of Aldolase knockdown —which would reduce glycolytic flux— and highlights the role of metabolic regulation within our co-expression module and its relevance to epilepsy.

### Deregulated energy metabolism as a common denominator of altered seizure susceptibility in the *Drosophila co-expression module*

Unbiased KEGG pathway and GO-term enrichment analyses revealed a contribution of carbohydrate metabolic processes to the fly co-expression module. Strikingly, we experimentally determined that altered energy metabolism (i.e., metabolic rate) was not at all limited to *Ald1* (the only viable condition of the three genes —*Ald1*, *Eno*, and *Pgi*— operating in carbohydrate metabolic processes) but was a highly penetrant feature among the pan-neuronal knockdown models with altered susceptibility to seizures. Of note, this included models for which the underlying gene has been previously connected to energy metabolism (*14-3-3ζ, CaMKII*, *NP15.*6, *Cam,* and *Vha68-*1; see **Table S8**) as well as genes with no previous indication for a regulatory function in energy metabolism (*shep* and *aSnap*; **Table S8**).

There is accumulating evidence for brain metabolic dysfunction, particularly hypermetabolism, contributing to epilepsy (Rho and Boison, 2022). Hypermetabolism is frequently considered a compensatory response, especially in situations where increased neuronal activity drives higher energy consumption (Ashraf et al., 2015, Liotta et al., 2024). A direct contribution of hypermetabolism to epilepsy is further supported by the clinical efficacy of ketogenic diet (D’andrea Meira et al., 2019), and anti-epileptic drugs such as the lactate dehydrogenase-inhibitor stiripentol (Sada et al., 2015). Furthermore, glycolysis inhibitors such as 2-deoxy d-glucose (2-DG) have been reported to reduce seizure susceptibility in several animal models and cellular systems (Shao et al., 2018). Most of our knockdown models also exhibited increased metabolic rates, leading us to propose that hypermetabolism, which is a common feature of OXPHOS deficiency (Sturm et al., 2023), plays a significant role in regulating seizure susceptibility.

However, there is also a body of literature reporting hypometabolism in epilepsy as well as pro-epileptic effects of glycolytic inhibition. The reasons for the observed discrepancies have been proposed to include acute versus chronic effects (Liotta et al., 2024, Samokhina et al., 2017, Stafstrom et al., 2009), different types of metabolic disorders (Bartolini et al., 2023, Klepper et al., 2020), variations in epileptogenic zones (Carvalho et al., 2022, Schur et al., 2018), differences between excitatory and inhibitory deficits (Bowser and Khakh, 2004, Chassoux et al., 2016), and positive feedback or compensatory effect of dynamic changes (Koenig and Dulla, 2018, Otahal et al., 2014), all of which can in part contribute to opposite effects. Thus, the relationships between brain metabolism and epileptic seizures are complex (Rho and Boison, 2022). This complexity may also be underlying our finding of hypermetabolism in both subgroups of our models, those with increased as well as with decreased susceptibility for seizure-like behavior. Further investigating the metabolic substrates that fuel increased metabolic rate in our models may hold some answers. Regardless of this knowledge gap, our study revealed a group of disorders that represent highly feasible candidates for metabo-therapeutic interventions.

### AMPK signaling in *Drosophila* models with altered seizure-like behavior

To ask whether the identified role of the co-expression module in the control of metabolic rate may be explained by convergence onto a specific molecular pathway, we turned to AMPK, an energy and glucose sensing protein (Lin and Hardie, 2018). Previous work has shown increased levels of p-AMPK upon chemically induced seizure in mice models (Lee et al., 2009). Consistent with this finding, our pan-neuronal knockdown models showing seizure-like behaviors lead to increased AMPK phosphorylation, though regardless of their metabolic rate (with p-AMPK also being increased in *cype* with unaltered and *Vha68-1* with decreased metabolic rate). To further investigate this surprising finding, we assessed additional controls. These revealed a modest but significant increase in AMPK phosphorylation upon RNAi targeting genes not associated with our 26 gene co-expression module as well as a control with RNAi against a non-*Drosophila* gene. Of note, this finding was specific to induced RNAi; we neither observe changes in p-AMPK levels in non-induced UAS-RNAi models or Gal4 only controls. This finding is alarming as it suggests a target-independent effect of the activated RNAi machinery on AMPK signaling, making it challenging to interpret some of our findings. However, p-AMPK levels in controls were considerably lower than in some of our pan-neuronal knockdown models (*14-3-3ζ*, *NP15.6*, *Ald1*, and particularly *Cam and CaMKII*). While these findings alone are not sufficient to explain the absence versus presence of seizures, nor the directionality of changes in metabolic rate, inducing expression of either wild-type or constitutively active AMPK construct conferred a dose-dependent protective effect against heat-induced seizures. This aligns with studies reporting that time-restricted feeding had an anticonvulsant effect and resulted in increased p-AMPK levels as well as with findings that metformin, an AMPK activator, shows potential anti-epileptic effects (Alnaaim et al., 2023, Landgrave-Gomez et al., 2016). Our results suggest that activated AMPK can alleviate seizure susceptibility in a dose-dependent manner. Expression of the dominant-negative AMPK allele, in contrast, had no effect on seizure susceptibility. Altogether, we propose that increased AMPK activation may underlie accelerated metabolism to alleviate seizure susceptibility in some of our models yet conclude that other mechanisms must contribute. Hence, our co-expression module appears to converge on altered metabolic rate, but not on a uniform underlying molecular mechanism.

### Limitations and future directions

Despite the discovery of metabolic dysregulation in a number of rare epilepsy disorder models linking synaptic and metabolic mechanisms, our study has some limitations. In addition to the complex relationship between metabolic rate, p-AMPK levels, and altered seizure susceptibility, further limitations are based on decisions that have guided our bioinformatics approach. Specifically, we refined the highly epilepsy-associated human M30 gene network using data from the first *Drosophila* scRNA-seq brain dataset (Davie et al., 2018), which, although pioneering, had less sequencing depth compared to more recent studies (Jovic et al., 2022, Li et al., 2022). We also decided to focus our study on co-expression in neurons, given that the core physiological feature of epileptic seizures is neuronal hyperexcitability (Holmes and Ben-Ari, 2001). However, an increasing number of studies provide compelling evidence for glial involvement in the pathophysiology of epilepsy (Patel et al., 2019). Further investigations into whole-brain or perhaps even glia-specific co-expression can be expected to contribute additional converging themes in epilepsy.

In conclusion, we have shown that single-cell expression data and experimental follow-up in *Drosophila* can refine human disease-relevant GCNs, highlighting the potential of integrating cross-species data to enhance our understanding of human diseases. Our work identified a co-expressed gene module, conserved across evolution and operating at the interface of synaptic and metabolic function, as a converging mechanism in epilepsy. We consider this particularly interesting considering emerging evidence for local glycolytic and mitochondrial mechanisms in nerve terminals/at synapses (Faria-Pereira and Morais, 2022, Li and Sheng, 2022, Myeong et al., 2023, Pulido and Ryan, 2021). Our findings that genetically disturbing synaptic key players (*CaMKII* and *Vha68-1*), reversely, lead to changes in metabolic rate indicates previously unappreciated mechanisms, crosstalk, or feedback mechanisms. To further understand these will provide fundamental insights that may open novel opportunities to interfere with intertwined synaptic and metabolic epileptic pathologies.

## MATERIALS AND METHODS

### Gene orthology and single-cell RNAseq co-expression analysis

The orthologs of human genes from the epilepsy-associated M30 network (Delahaye-Duriez et al., 2016) were identified using FlyBase (release FB2020_02) (Gramates et al., 2022, Jenkins et al., 2022) with DIOPT scores more than or equal to three. 163 M30 genes were identified as one-to-one fly orthologs. In addition, 37 M30 genes represented members of shared, conserved gene families with reduced redundancy in the fly, leading to the inclusion of 17 many-to-one *Drosophila* orthologs. Lastly, 31 M30 genes had more than one single *Drosophila* homolog with high and similar DIOPT scores, leading to the inclusion of 107 one-to-many fly genes, together resulting in a catalog of 287 unique M30-related *Drosophila* genes. Single-cell RNA sequencing (scRNA-seq) data from whole-brain Drosophila neurons were obtained from a previously published (Davie et al., 2018, Hwang et al., 2018). The dataset “Aerts_Fly_AdultBrain_Filtered_57k.loom” was downloaded from https://scope.aertslab.org/#/Davie_et_al_Cell_2018/, and the expression matrix and the cell type annotations were extracted using the SCopeLoomR package. Of the 287 M30-related *Drosophila* genes, 142 were retained after requiring a minimum expression of 5000 reads across all cells in the dataset. To focus on relevant neuronal populations, a coherent dataset of defined neuronal cell types representing diverse neuronal functions was selected, including populations of serotonergic, tyraminergic, peptidergic, dopaminergic and octopaminergic neurons (adPN, adPN/C15, adPN/C15&kn, adPN/kn, adPN/kn&CG31676, AstA/NPF, AstA/Nplp1, Capa, CCAP, CCHa1, Clock, Crz, DCN, DN1, Dopaminergic, dorsal_Fan-shaped_Body, FMRFa, Gr43a, Hsp, Hug, ITP, L1, L2, L3, L4/L5, Lamina_monopolar, Lawf1, Lawf2, LNv, lPN, lPN/CG31676, lPN/unpg, MBON, Mip, Mip/ITP, Mip/OCT, Octopaminergic, Olfactory_projection_neurons, PAM, Peptidergic, Poxn, Proc/Gpb5, Proc/Ms, Serotonergic, Tyraminergic). This selection excluded cell types that had previously been reported to possess strongly distinct transcriptomics profiles as they could bias the correlation analysis (glial cells, optic lobe neurons, photoreceptors and Kenyon cells (Davie et al., 2018, Hwang et al., 2018). Gene co-expression relationships were evaluated by calculating pairwise Spearman’s correlation coefficients across all selected neuronal cells (i.e. correlation on the 142 genes x 5980 cells log2-transformed matrix). Correlation values were assembled into a gene-gene correlation matrix. Genes were hierarchically clustered based on pairwise correlation distances to identify co-expression modules (hclust(as.dist(1-geneCor)) and visualized using the NMF::pheatmap function (Kolde, 2019).

### Classification of epilepsy-associated genes and statistical analysis

We divided human orthologs of genes in the fly co-expression module into three categories: epilepsy genes, epilepsy-related genes and genes not associated with epilepsy to date. For this, human orthologs were first searched in the OMIM database (<u>OMIM</u>). Genes with mentioning of epilepsy or an epilepsy-related term (such as epileptic encephalopathy) in the main phenotype description field of the Gene-Phenotype Relationships table were categorized as “epilepsy gene”. Genes for which epilepsy or seizures were listed in OMIM’s clinical features section or for which we found case reports in the literature, retrieved by search terms “epilepsy” or “seizure”, were categorized as “epilepsy-related gene”. Genes that did not fall into the epilepsy or epilepsy-related categories were classified as not associated with epilepsy. Statistical analysis of enrichment for epilepsy-associated genes (comprising epilepsy and epilepsy-related genes) was performed using cumulative distribution function of hypergeometric distribution with the <u>Hypergeometric *p*-value calculator (ucla.edu)</u>.

### Gene Ontology (GO) and Kyoto Encyclopedia of Genes and Genomes (KEGG) pathway analyses

GO for biological processes (GO: BP) and KEGG analysis was performed for the bipartite fly co-expression module (n = 26) with all *Drosophila* genes used as the background, as well as for the human M30 cluster (n = 320) with all human genes as the background, using g:Profiler (Raudvere et al., 2019). Terms were defined as significant if the Benjamini-Hochberg FDR was < 0.05. A selection of significant GO: BP terms were visualized using a dot-heatmap generated using *ggplot2* to display both FDR significance and enrichment (Wickham, 2016). The complete list of significantly enriched GO: BP and KEGG terms is provided in **Table S5**. Hypergeometric test statistics and overlap between significant GO terms in the fly co-expression module and human M30 cluster was determined in R (R Core Team, 2021) using a population size of all potential GO: BP terms (27,047), and visualized using a Venn diagram generated using BioVenn (Hulsen et al., 2008).

### *Drosophila* stocks & maintenance

Flies were cultured in a standard medium containing cornmeal, yeast, sugar, and agar. The rearing conditions were maintained at a constant temperature of 25°C and 60% relative humidity, under a 12:12-hour light-dark (LD) cycle. The Gal4/UAS system was used to study the effect of neuronal-specific knockdown of different genes (Brand and Perrimon, 1993). For this purpose, the pan-neuronal driver *nSyb-Gal4* [Bloomington *Drosophila* Stock Center (BDSC) #51635] was crossed to UAS-RNAi lines. UAS-RNAi lines were obtained from the VDRC or BDSC and are summarized in **Table S7**. Further *Drosophila* stocks that were used in this study are: the genetic background controls of the KK library [Vienna *Drosophila* Resource Center (VDRC) #60100], of the GD library (VDRC #60000), and of the TRiP libraries attP2 (BDSC #36303), and attP40 (BDSC #36304). To generate the experimental control animals, lines containing the corresponding genetic background to the RNAi constructs (VDRC #60000, VDRC #60100, BDSC#36303, BDSC#36304) were crossed to the driver line.

### Mechanical induced seizure-like behavior

The mechanically-induced seizure-like behavior assay was adapted from Kuebler and Tanouye (Kuebler and Tanouye, 2000). At day of eclosion (day 0), five male flies per genotype were collected under CO_2_ anesthesia and housed in vials with standard food. Animals were transferred to fresh food on day three. At six days old, flies were allowed to acclimate for five minutes in new, transparent vials and were sequentially mechanically stimulated in a vortex mixer at the maximum setting (2400 RPM) for one minute, and videos were recorded after the stimuli. The number of flies that showed seizure-like behavior [either lying on their backs with uncontrolled wing flapping and/or leg twisting (Kuebler and Tanouye, 2000) within five seconds or paralysis for more than five seconds post-stimuli] was scored. The entire procedure was conducted under double-blind conditions to ensure unbiased scoring, and the genotype identities were only decoded post-analysis. Statistical analyses were performed using GraphPad Prism version 10.1.2 for Windows (GraphPad Software, San Diego, California, USA). Since the dataset did not follow a Gaussian distribution, it was analyzed using Kruskal-Wallis test with Dunn’s multiple comparisons test. Significance in the figures correspond to by adjusted *p*-values, reflecting only those results meeting the corrected significance threshold. All data presented were derived from two (Figure 3) or three (Figure 6) independent experiments.

### Heat induced seizure-like behavior

The heat induced seizure-like behavior assay was adapted from Fischer and colleagues (Fischer et al., 2016). Flies were reared at 25°C or 28°C as indicated. For quantifying heat induced seizure-like behavior, aliquots of five males per genotype were collected at zero days post-eclosion under CO_2_ anesthesia, and housed in vials with standard food, with a transfer to fresh vials on day three. At six days old, flies were acclimated for five minutes in new, transparent vials. Vials were individually submerged in a temperature-controlled water bath with and adjusted to the target temperature with a tolerance of ± 0.2 °C. Flies in the KK and TRiP genetic backgrounds were immersed at a temperature of 37°C, while those in the GD background were exposed to 38°C based on their different temperature sensitivity. The number of flies showing seizure-like behavior was recorded at five-minute intervals over a duration of 80 minutes. For testing the ability of AMPK overexpression under stringent conditions, flies were immersed into a 42°C water bath. The number of flies showing seizure-like behavior was recorded at 10 seconds intervals over a duration of 10 minutes. To calculate the time at which 50% of individuals showed seizures, a sigmoidal curve with a variable slope was applied to the raw data. This calculation of the time per genotype, referred to as logEC50, was performed using GraphPad Prism. This timepoint serves as an indicator of seizure susceptibility and is utilized for subsequent statistical analyzes. The values for the RNAi and overexpression lines were normalized against their respective background controls, due to day-to-day variation. The entire procedure was conducted under double-blind conditions to ensure unbiased scoring, and the genotype identities were only decoded post-analysis. Statistical analyses were performed using GraphPad Prism. Since the dataset was normally distributed, it was analyzed using ordinary one-way ANOVA tests with Šídák’s multiple comparisons test. Significance in the figures is denoted as adjusted *p*-values, reflecting only those results meeting the corrected significance threshold. All data presented were derived from two (Figure 3) or three (Figure 6) independent experiments.

### Metabolic rate

The metabolic rate assay was adapted from Yatsenko and colleagues (Yatsenko et al., 2014). In brief, to measure CO_2_ production in *Drosophila*, aliquots of 25 males per genotype were collected after eclosion (day 0-1) under CO_2_ anesthesia, gently transferred into vials containing standard food, and flipped to new vials on day three. For behavioral testing, 6–7-day old flies were transferred without CO_2_ anesthesia to homemade respirometers in groups of five and allowed to acclimate for 15 minutes.

The respirometers, crafted using pipette tips and capillary micropipettes with soda lime as a CO_2_ absorbent, were hung vertically to immerse the tip into a bromophenol blue solution, and the chambers were sealed. Following a 75-minute acclimation period within the sealed chamber, photos were taken documenting the liquid level at the start and end of a 1-hour period. Photos were analyzed using Fiji 1.53 to measure the liquid’s travel distance. The resulting data were presented as microliters of CO_2_ produced per fly. Statistical analyses were performed using GraphPad Prism. Datasets following a Gaussian distribution were analyzed with two-tailed unpaired t-tests. Datasets that did not follow a Gaussian distribution were analyzed using Mann-Whitney tests. All data presented were derived from three independent experiments.

### Novelty of identified links between genes with altered seizure susceptibility, energy metabolism and AMPK activity

To provide an overview on the literature implicating genes of the evolutionarily conserved and co-expressed 26-gene module with altered seizure susceptibility in energy metabolism and/or AMPK activity, a comprehensive search of the PubMed database was conducted for each gene using terms “metabolic rate”, “AMPK”, “CO_2_ production”, “O_2_ consumption”, “energy metabolism” and “glycolytic flux”. Relevant matches are listed in **Table S8**.

### Western blotting

To detect AMPK phosphorylation levels aliquots of 25 males per genotype were collected after eclosion (day 0-1) under CO_2_ anesthesia. They were gently transferred into vials containing standard food and subsequently flipped to new vials on day 3. After six days, flies were CO2 anesthetized, transferred to Eppendorf tubes and snap-frozen in liquid nitrogen. Heads were separated from bodies by flicking the tubes and were collected with a brush at 4°C. Twenty heads per sample were homogenized in 80 μl RIPA buffer with phosphatase inhibitors. Samples were mixed with a 1:1 volume of NuPAGE LDS Sample Buffer 4x (Life Technologies, cat# NP0007, Inchinnan, UK) with DL-Dithiothreitol solution (Sigma-Aldrich, cat# 43816-10ML, Saint Louis, US), and boiled for 10 minutes. 12 μL of each sample was loaded onto a 15-well NuPAGE 4-12% Bis-Tris gel (Life Technologies, cat# NP0323BOX, Inchinnan, UK) and electrophoresed for 15 minutes at 80V, followed by 120 minutes at 120V. Proteins were transferred to nitrocellulose membranes (Trans-Blot Turbo Transfer Pack, Bio-Rad, cat# 1704158, Hercules, CA) using the Trans-Blot Turbo Transfer System (Bio-Rad, cat# 1704150, Hercules, CA) at 25V for 7 minutes. Membranes were blocked for one hour in 10% bovine serum albumin (BSA, Sigma-Aldrich, cat# A8806-1G, Saint Louis, US), and then incubated in primary antibody diluted in a 1:1 volume of 10% BSA and tris buffered saline buffer with 0.2% tween (TBST) overnight at 4°C. The antibody against the phosphorylated form of α-AMPK (Cell Signaling, cat# 2535S, Danvers, MA; RRID:AB_331250) was diluted 1:500; anti-β-Tubulin (DSHB, cat# E7, Iowa, US; RRID:AB_528499) was diluted 1:6000. The membranes were then washed three times for 10 minutes each in TBST. Membranes were incubated in a goat anti-mouse IgG (H+L) secondary antibody (dilution 1:5000, Thermo Fisher Scientific, cat# A-21057, Waltham, US; RRID: AB_2535723) and a goat anti-rabbit secondary antibody (dilution 1:5000, Li-cor, cat# 926-32211, Lincoln, NE; RRID:AB_621843), diluted in a 1:1 volume of 10% BSA and TBST buffer at room temperature, for two hours in the dark. Three 10-minute washes with TBST buffer were done and immersed in tris buffered saline in the dark. Membranes imaged on an Odyssey Infrared Scanner (Odyssey® DLx Imaging System, Li-cor, cat# 9142-00, Lincoln, NE). Quantification was conducted using Image Studio software (version 4.0.21). All direct comparisons were only done on samples run on the same gel, and the Odyssey Integrated Intensity was used to estimate the relative strength of signals across different blots. Each experimental group comprised three biological replicates from two independent experiments (six samples), and each sample run once on a gel for quantification. Statistical analyses were performed using GraphPad Prism. Datasets following a Gaussian distribution were analyzed with two-tailed unpaired t-tests. All data presented were derived from two independent experiments.

## Acknowledgements

We acknowledge Stein Aerts and the Aerts laboratory for their resources and support with the single-cell RNA sequencing data. We thank the VDRC and the Bloomington *Drosophila* Stock Center for reagents. We are grateful to I. Janssen, J. Luyckx, L. Gerding, M. de Wit, S. Letteboer, and S. Beersum for experimental or logistic support, and to all members of the Schenck lab for helpful discussions.

## Footnotes

### Author contributions statement

Conceptualization: J.L., K.L., M.C.-T., A.Schenck, Methodology: J.L., S.G.J., B.R., F.K., S.A., M.A.H., Software: S.G.J., S.A., M.A.H., Formal analysis: J.L., S.G.J., A.Serna, K.L., Investigation: J.L., S.G.J., A.Serna, Data curation: J.L., S.G.J., Visualization: J.L., S.G.J., M.C.-T., A.Schenck, Supervision: P.V., K.L., M.C.-T., A.Schenck, Project administration: J.L., M.C.-T., A.Schenck, Funding acquisition: J.L., S.G.J., K.L., A.Schenck, Writing - original draft: J.L., M.C.-T., A.Schenck, Writing – review & editing: J.L., S.G.J., A.Serna, B.R., F.K., S.A., P.V., M.A.H., K.L., M.C.-T., A.Schenck.

## Funding

This work was in part supported by a personal fellowship from the China Scholarship Council (201908440286) to J.L., by Radboud Excellence Fellowships to S.G.J. and K.L., and by a Vici grant from the Netherlands Organization for Health Research and Development (ZonMw, 09150181910022) to A.S..

## Data and resource availability

All relevant data and resources can be found within the article and its supplementary information.

## Competing interests

The authors declare that they have no competing interests.

## Notes

### Competing Interest Statement

The authors have declared no competing interest.

